# Taming the data deluge: a novel end-to-end deep learning system for classifying marine biological and environmental images

**DOI:** 10.1101/2022.10.20.509848

**Authors:** Hongsheng Bi, Yunhao Cheng, Xuemin Cheng, Mark C. Benfield, David G. Kimmel, Haiyong Zheng, Sabrina Groves, Kezhen Ying

## Abstract

Marine underwater imaging facilitates non-destructive sampling of species at frequencies, durations, and accuracies that are unattainable by conventional sampling methods. These systems necessitate complex automated processes to identify organisms efficiently, however, current frameworks struggle to disentangle ecological foreground components from their dispensable background content. Underwater image processing relies on common architecture: namely image binarization for segmenting potential targets, prior to information extraction and classification by deep learning models. While intuitive, this infrastructure underperforms as it has difficulty in handling: high concentrations of biotic and abiotic particles, rapid changes in dominant taxa, and target sizes that vary by several orders of magnitude. To overcome these issues, a new framework is presented that begins with a scene classifier to capture large within-image variation, such as disparities in particle concentration and dominant taxa. Following scene classification, scene-specific regional convolutional neural network (Mask R-CNN) models were trained to separate target objects into different taxonomic groups. The procedure allows information to be extracted from different image types, while minimizing potential bias for commonly occurring features. Using *in situ* coastal PlanktonScope images, we compared the scene-specific models to the Mask R-CNN model including all scene categories without scene classification, defined as the full model, and found that the scene-specific approach outperformed the full model with >20% accuracy in noisy images. The full model missed up to 78% of the dominant taxonomic groups, such as *Lyngbya, Noctiluca*, and *Phaeocystis* colonies. This performance improvement is due to the scene classifier, which reduces the variation among images and allows an improved match between the observed taxonomic groups and the taxonomic groups in pre-trained models. We further tested the framework on images from a benthic video camera and an imaging sonar system. Results demonstrate that the procedure is applicable to different types of underwater images and achieves significantly more accurate results than the full model. Given that the unified framework is neither instrument nor ecosystem-specific, the proposed model facilitates deployment throughout the marine biome.

## I. Introduction

Imaging systems are increasingly being used to study aquatic organisms and their interactions with the environment at different spatial and temporal scales [1-6]. Despite the rapid developments and applications of various imaging systems, automated image processing faces several challenges: The root of the problem lies in in the field’s lack of an adopted unified processing framework. Most automated image processing procedures are customized and often system-specific[1, 7], a fact that hinders broader application. This phenomenon is exacerbated by environment-driven separation protocols, which can require varying levels of disassociation of target and non-target objects and separation of background and foreground objects. Further challenges include: segmentation method, unbalanced taxonomic samples, size class differences, particle saturation, overcrowding, and overlap.

The challenges of developing a general image processing framework arise from differences in image properties associated with different imaging technologies and different environments. For example, processing plankton images primarily means separating target and non-target objects in the water column, whereas processing benthic images requires separation of foreground objects from both the background and non-target objects. In previous cases, these tasks have been distinguished and processed by convolutional neural networks (CNNs) [8-11]. A commonly applied architecture in aquatic biological images starts with potential targets, denoted by the region of interest (RoI), and the segmented RoIs are standardized before feature description and classification by a pretrained machine learning model [6, 8, 12, 13]. However, accurate segmentation and class unbalance remain two major issues, both of which undermine the performance and accuracy of the common framework by producing excessive false positives and biased results.

The goal of accurate segmentation of RoIs from underwater images is to separate individual organisms and ensure that each ROI only has one potential target for the subsequent classification. While accurate segmentation is relatively easy for images collected in water with low particle density, it remains a challenge in water with higher particle density (Fig. 1). Using plankton images collected from coastal waters as an example, images are often crowded either with nonbiological particles or planktonic organisms (Figs.1a -c). When images are saturated with nonbiological particles, it is difficult to separate target organisms from non-target particles (Fig 1a). When organisms have a complex structure, over-segmentation often occurs [12]. Furthermore, overlap among different organisms makes the task of segmenting individuals even more challenging (Figs 1 a-e). The issue of crowded images and overlap among different organisms arises from two different aspects. Patchy distribution is a common feature of marine organisms [14], and when an imaging system is towed through a patch of marine organisms, we would expect overcrowded images and overlap among individual organisms. As technology advances, modern imaging systems are often equipped with increased field of view and depth of field, which further exacerbate the issue of overcrowded and overlapping objects.

**Fig. 1.**
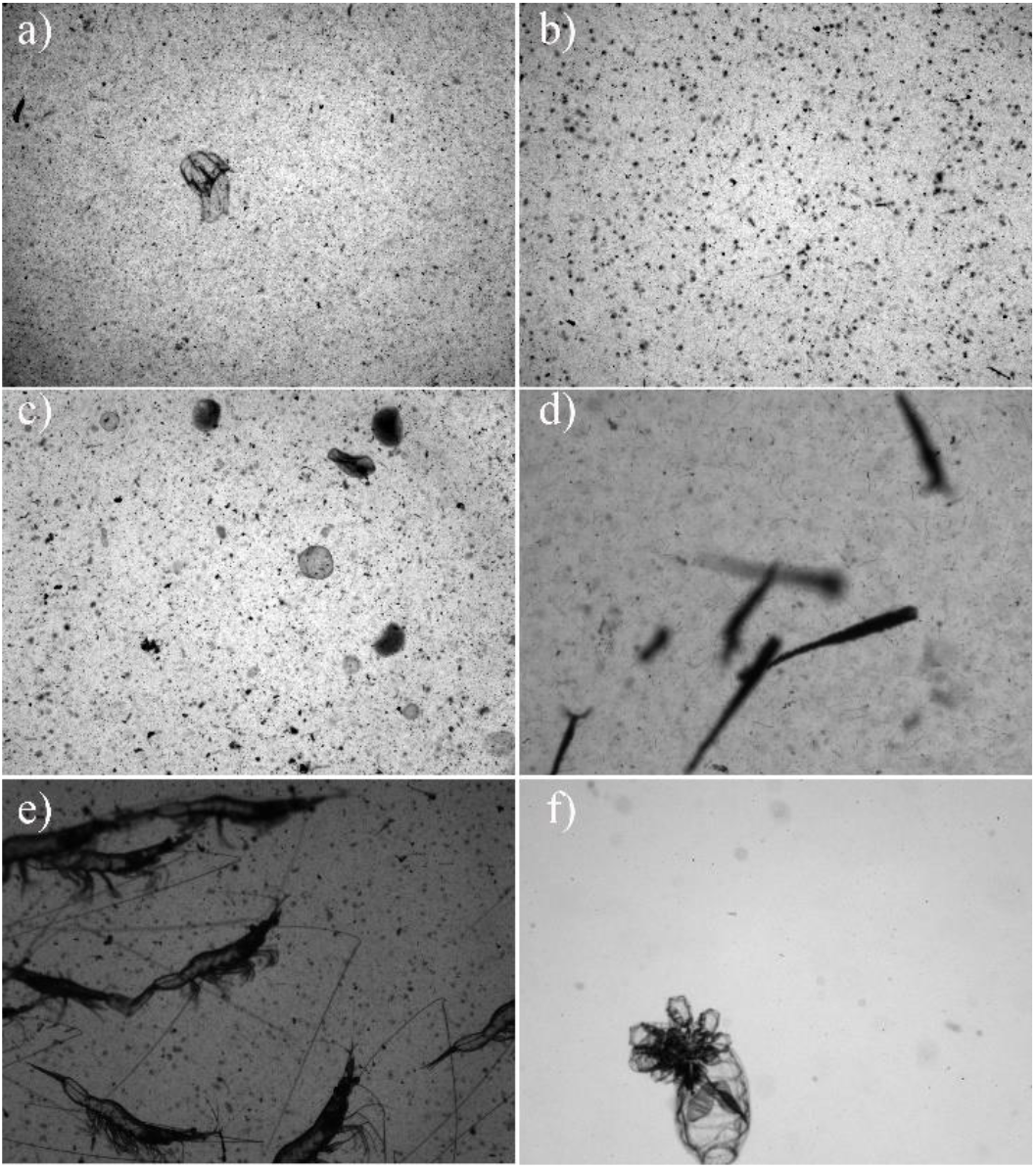
Example images for selected scene categories: a) High concentration scene with large amounts of nonbiological particles, b) *Noctiluca* scene with each black dot indicating one *Noctiluca*, c) *Phaeocystis* scene showing the semi-transparent round-shaped colonies, d) Pteropoda scene showing clustered *Creseis acicula* and line-shaped *Lyngbya* algae, f) Shrimp scene showing clustered individuals, and g) Low density scene showing a clean image with one budding Thaliacea.

Image segmentation is a long-standing problem in computer vision and most existing techniques are not suitable for noisy environments [15, 16]. Recent efforts on developing segmentation techniques, specifically for crowded underwater images, partially alleviate this issue [17, 18]; but given the complexity and uncertainty in underwater images, unsupervised deep learning approaches like region-based CNN (R-CNN) offer a more promising solution [19]. The R-CNN models generally include a region proposal network (RPN) to locate RoIs, a CNN model to describe features of RoIs generated from RPN proposals, and a classification layer to predict final bounding boxes and classes. Mask R-CNN is a combination of Faster R-CNN and a fully convolution network which outperforms existing models in instance-level segmentation and recognition [20, 21]. In the present study, we test the feasibility of using Mask R-CNN as the first step in processing noisy marine biological and environmental images.

In marine environments, skewed frequency distribution across taxonomic groups is common and such imbalanced class distributions have detrimental impacts on classification performance because the model often oversamples the rare groups and under samples the abundant groups [22-24]. While a more balanced class distribution fits our preference for unbiased results for both common and rare taxonomic groups, deep learning models require tens of thousands of labelled images to achieve high accuracy, which is almost impossible for rare taxonomic groups. Meanwhile, the large size variation that spans up to several orders of magnitude, also complicates the imbalance issue. Large organisms occupy the greatest numbers of pixels, and morphological features can persist throughout the convolution process; whereas small organisms occupy small amounts of pixels and morphological features likely disappear after a few iterations of convolution. Therefore, small organisms need more labelled images than large organisms to reach the same level of accuracy. This training imbalance can either exacerbate or alleviate the unbalance class issue, depending on the steps that precede convolution.

There is no simple solution to the common issue of unbalanced classes in deep learning, but proper data-level operations could be helpful in addressing this issue [24]. We propose to alleviate the issue of unbalanced class at the data level by incorporating a scene classification model. Scene classification uses the layout of organisms within the scene, in addition to the ambient context, to group similar images. Using underwater plankton images as an example, images with high concentration of nonbiological particles (Fig. 1a) are often collected around locations or times with strong physical mixing. Single-species dominated images are often collected within a bloom period or a patch of the organisms (Figs 1c-e). Scene classification reduces the diversity and uncertainty within each category, allowing a more targeted classification model to be trained. It also facilitates better matching between observed frequency across different taxa and the frequency distribution within the model’s library. For example, images from clear water are often characterized by low concentration of biological and nonbiological particles (Fig. 1f) with a less skewed observed frequency across different taxa than images collected within a bloom or patch, therefore, a library built from these clear water images naturally provides a better match.

In the present study, we propose a general framework to process underwater images (Fig. 2). The procedure starts with scene classification to separate images based on their context and layout of dominant organisms. This process is followed by a separate Mask R-CNN model, trained for each scene, to facilitate foreground object detection and recognition. To determine how the new framework compares to prior methods, we provide two models: full model, scene-specific model. The full model represents the common architecture, and is accomplished through a Mask R-CNN model for automated recognition of underwater organisms without scene classification. The scene-specific model starts with a scene classification model followed by a dedicated model for each scene for recognition. We provide the metrics for each approach using underwater plankton images from coastal waters collected by PlanktonScope [25, 26]. We then expand the comparison to test for general applicability using images collected by different underwater imaging systems.

**Fig. 2.**
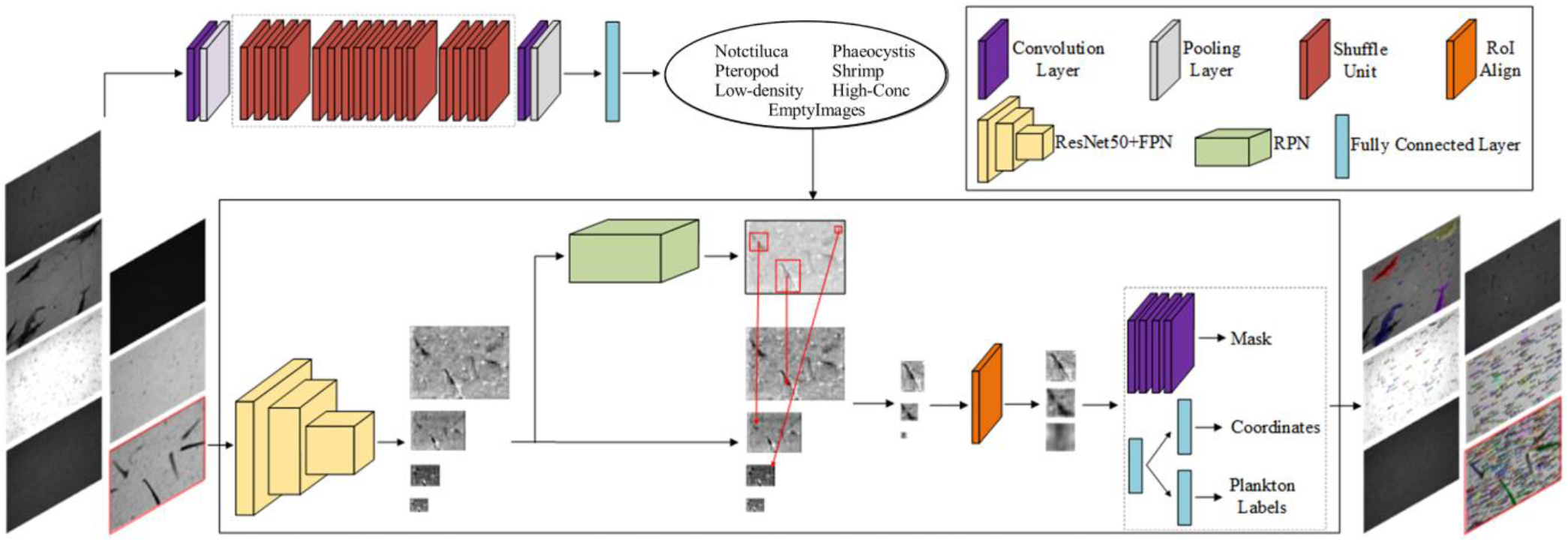
Diagram of the proposed procedure. The procedure starts with a scene classification using a lightweight ShuffleNet. For example, six scenes and an additional scene for empty images were selected based on image contents such as dominant taxa group or concentration of particulates. A separate Mask Region-Based Convolutional Neural Network (RCNN) model was trained for each scene category and potential objects were first detected through a region proposal network and then classified by a residual neural network (lower box). The final output included an image with predicted results and segmented objects for each taxa group.

## II. Methods

### A. Full model

Mask R-CNN combines a Faster CNN for object identification and a fully convolutional network for recognition. Faster CNN includes two networks, a CNN and RPN. First, the network detects region proposals that are defined as regions in the feature map which contain the object. In the second stage, the network predicts bounding boxes and object class for each of the proposed regions obtained in the first stage. Each proposed region can be of different size and the size of these proposed regions is then fixed by the RoI pooling method, which helps to preserve spatial information. Mask R-CNN extends Faster R-CNN by adding a branch for predicting an object mask in parallel with the existing branch for bounding box recognition. It also replaces the RoI pooling with RoIAlign which makes object detection more efficient and accurate while simultaneously generating a high-quality segmentation mask for each instance. The output from the RoIAlign layer is analyzed by a residual neural network (ResNet) for object classification. Finally, the procedure segments recognized objects based on the corresponding mask generated during object detection [20].

The Mask R-CNN model was implemented in Python using the Detectron 2 package [27]. In the present study, 960 images labelled using the open source annotation tool LabelMe [28]. Within these images, we identified 17 taxonomic groups and labelled 32, 227 individual organisms ranging from small algae, <100 µm in diameter, to small pelagic fish, ∼4 cm. The first three abundant taxonomic groups are the line-shaped algae *Lyngbya*, near-transparent ellipsoid *Noctiluca*, and copepods (Table I). We trained a full model, using the labelled images, and assessed the accuracy of foreground object identification and classification to evaluate the model performance.

**TABLE I.** Number of Images, Identified Taxonomic groups, AND Labelled Individuals for Each Taxonomic Group in the Full Model and Scene-specific Models.

|  | Full model | Different scenes |  |  |  |  |  |
| --- | --- | --- | --- | --- | --- | --- | --- |
|  |  | High Concentration | Noctiluca | Phaeocystis | Pteropod | Shrimp | Low density |
| Number of images | 970 | 151 | 99 | 84 | 324 | 124 | 431 |
| Appendicularia | 489 | 29 | 13 | 147 | 207 | 4 | 239 |
| Chaetognatha | 209 | 22 | 42 | 45 | 12 | 9 | 81 |
| Copepoda | 3,466 | 144 | 353 | 531 | 1,039 | 119 | 1,952 |
| *Creseis* | 1,225 | 0 | 0 | 0 | 1,225 | 9 | 188 |
| Echinodermata | 451 | 5 | 388 | 34 | 16 | 0 | 12 |
| Fish | 10 | 0 | 0 | 0 | 0 | 10 | 0 |
| Larval Fish | 1 | 0 | 0 | 0 | 0 | 0 | 3 |
| *Lyngbya* | 20,212 | 1,635 | 1,593 | 815 | 1,8370 | 205 | 10,107 |
| Medusae | 246 | 73 | 28 | 38 | 50 | 6 | 109 |
| *Noctiluca* | 4,950 | 189 | 3,661 | 4 | 6 | 782 | 595 |
| *Phaeocystis* | 521 | 143 | 86 | 259 | 40 | 0 | 0 |
| Polychaete | 21 | 0 | 0 | 0 | 0 | 0 | 21 |
| *Scylla* larvae | 14 | 0 | 0 | 0 | 0 | 0 | 14 |
| Shrimp | 262 | 76 | 7 | 0 | 2 | 154 | 7 |
| *Skeletonema* | 3 | 2 | 0 | 0 | 1 | 0 | 0 |
| Spiral_diatom | 111 | 13 | 16 | 0 | 24 | 0 | 15 |
| Thaliacea | 108 | 9 | 54 | 5 | 13 | 5 | 57 |

### B. Scene-specific model

The scene-specific model begins by classifying underwater images into different scenes, and is followed by a scene-specific Mask R-CNN model for objection detection and classification (Figure 2). Scene classification depends on the ambient context, shape, and layout of biological targets within the image. This step is accomplished through an efficient lightweight CNN model, namely ShuffleNet, which utilizes pointwise group convolution and channel shuffle to reduce computation cost while maintaining accuracy [29, 30]. We use underwater plankton images to illustrate how we set up and train the scene classification model.

In the present study, we separate underwater plankton images into six scene categories (Fig. 1):

1. images with evenly distributed, abundant nonbiological particle (Fig. 1a), which are common in estuaries and nearshore waters due to riverine and resuspended bottom particles;
2. randomly distributed, *Noctiluca*-dominated images (Fig. 1b);
3. randomly distributed *Phaeocystis* colonies, which exhibit round clustering (Fig. 1c);
4. *Creseis acicula* Pteropoda clusters (Fig. 1d) and the co-occurring, evenly distributed, line-shaped *Lyngbya*
5. clustered, small shrimp (Fig. 1e) with co-occurring, randomly distributed *Noctiluca*;
6. clear water, low concentration plankton (Fig. 1f).

In the present study, we included 546, 1060, 768, 508, 336, and 766 full frame images in the high-concentration, *Noctiluca, Phaeocystis, Creseis*, shrimp, and clear water scenes respectively.

In the second phase, a separate Mask R-CNN model was trained for each scene category to detect and classify target objects (Fig. 2). Note that some images are intentionally included in different scenes repeatedly because most taxonomic groups occur in multiple scenes and the accuracy of scene classification is not 100%. For example, shrimp occur not only in the shrimp scene, but also in the high-concentration and low-density scenes. Meanwhile, some taxonomic groups often co-occur, and images could be separated into different scenes. For example, *Noctiluca* and shrimp often occurred together, and images could be classified within the *Noctiluca* scene or shrimp scene depending on the dominant group and the layout within the image. In both scenes, the co-occurring taxonomic groups are labelled. The number of images, taxonomic groups, and labelled targets for each scene were summarized in Table I. The number of images cross-referenced in different scenes were summarized in Table II.

**TABLE II.** Number of Images in Each Scene AND Cross-referenced Images Among Different Scenes.

|  | High-conc. | Noctiluca | Phaeocystis | Pteropoda | Shrimp | Low-den. |
| --- | --- | --- | --- | --- | --- | --- |
| High-Conc | 102 | 1 | 4 | 2 | 21 | 21 |
| Noctiluca | 1 | 85 | 3 | 3 | 2 | 5 |
| Phaeocystis | 0 | 1 | 76 | 0 | 0 | 4 |
| Pteropoda | 0 | 0 | 0 | 322 | 0 | 2 |
| Shrimp | 0 | 8 | 0 | 3 | 113 | 0 |
| Low-den. | 0 | 0 | 0 | 0 | 3 | 281 |

### C. Comparison between full model and scene-specific model

The performance of full model and scene-specific models was evaluated using accuracy metrics and a confusion matrix. During the process of model training, two accuracy metrics were calculated for the Mask R-CNN model: the accuracy for foreground object detection and the accuracy for object classification. For the scene classification model, a single accuracy metric was calculated. The general form of accuracy is 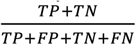, where TP is the number of true positives, TN is the number of true negatives, FP is the number of false positives, and FN is the number of false negatives. A confusion matrix was also constructed for the scene classification model. The confusion matrix includes the actual and predicted classes obtained by a classification system. Each row represents an actual class example; each column represents the state of a predicted class.

The performance of the full model and scene classification model were also evaluated using an underwater plankton dataset. The underwater plankton images were collected with PlanktonScope, an *in situ* shadowgraph imaging system in the coastal waters of Guangdong, P. R. China [25, 26] and the Columbia River Plume, U.S.A. Twenty full-frame images were selected for each scene, and then images were processed using both the full- and scene-specific models. Both models output the predicted results, segmented objects for each class, and the counts for each class. We then manually examined the segmented objects, moved the misclassified objects to the correct class, and performed a recount. Given that Mask R-CNN does not produce information for the non-target class and it is almost impossible to perform objective visual counts for small semi-transparent organisms like *Noctiluca* and *Lyngbya*, we used precision as a metric to evaluate model performance instead of accuracy. Precision was calculated as 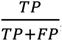, where TP is the number of true positive and FP is the number of false positive.

### D. General application

Benthic videos were collected in the Arctic using a camera system manufactured by A.G.O. Environmental Electronics Ltd., Victoria, B.C., Canada [31]. The system includes two positioning lasers, an undersea video camera, with onboard monitoring and recording on a ship-based video camcorder, and hand deployment using a 200 m electronic cable. A key challenge in processing benthic images is separating organisms from a complex background because benthic organisms often blend into their habitats and suboptimal image quality only exacerbates this problem. We compared the results of RoI extraction between the thresholding binarization approach and our approach. We separated benthic images into four scenes: images with aggregated organisms, complex background including dead shells and debris, isolated organisms, empty images without organisms. A scene classifier was trained with 120 images in each scene category. For demonstration purposes, we only included relatively few labelled images, 4 – 30, for each scene to train the scene-specific model. An independent set of 20 images were selected to test the performance of the model.

Adaptive resolution imaging sonar (ARIS) systems are ideal tools to study pelagic organisms ranging from jellyfish to small fish [32, 33]. Sonar images were collected in Chesapeake Bay using the ARIS 1800 (Sound Metrics Corp, Bellevue, WA, USA). These cameras have improved resolution and produce near-video quality images of the water column up to a 35 m range from the camera lens, and can resolve objects down to 3 mm. However, sonar images have much lower resolution than optical cameras and individual objects including small pelagic fish have few distinct morphological features. In most cases, visual identification relies on auxiliary information, such as schooling shape, size, and other spatial characteristics. For demonstration purposes, we separated sonar images into eight scenes: 1) empty images with only beam patterns; 2) sea floor; 3) fish schools without sea floor; 4) fish school with sea floor; 5) jellyfish without sea floor; 6) jellyfish with sea floor; 7) mysid shrimp swarms with sea floor; and 8) mysid shrimp swarms without sea floor. Note that the sea floor often has a strong signal and is an important feature to separate scenes because it both determines the layout of an image, and also affects image contrast. We also used an independent set of 20 images, with different organisms, to demonstrate the performance of the proposed framework.

## III. Results

### A. Comparison between full model and scene-specific model

The full model achieved 73% accuracy in foreground object detection and 87% accuracy in object classification using the labelled underwater plankton dataset (Fig. 3a). In the proposed framework, the accuracy of the initial step, scene classification, reached 98% using the selected full-frame plankton images (Fig. 3b) and the 6 scene-specific models generally reached higher accuracy than the full model (Figs. 3c-h). The confusion matrix (Fig. 4) suggests that the scene classification model can separate full frames into the corresponding scene with 94-99% accuracy for most scene categories. The frequent co-occurrence of shrimp and *Noctiluca* likely caused the relatively low accuracy in the shrimp scene. Among the scene-specific models, the high-concentration and shrimp-specific models had the highest accuracies, with 98% and 96% foreground classification accuracy and 97% and 96% classification accuracy respectively (Figs 3c, g). Pteropoda and low-density specific models had the lowest accuracies with 80% and 81% foreground classification accuracy and 89% and 92% classification accuracy. The increased scene-specific model performance suggests that scene classification reduces the variation, or uncertainty, among images.

**Fig. 3.**
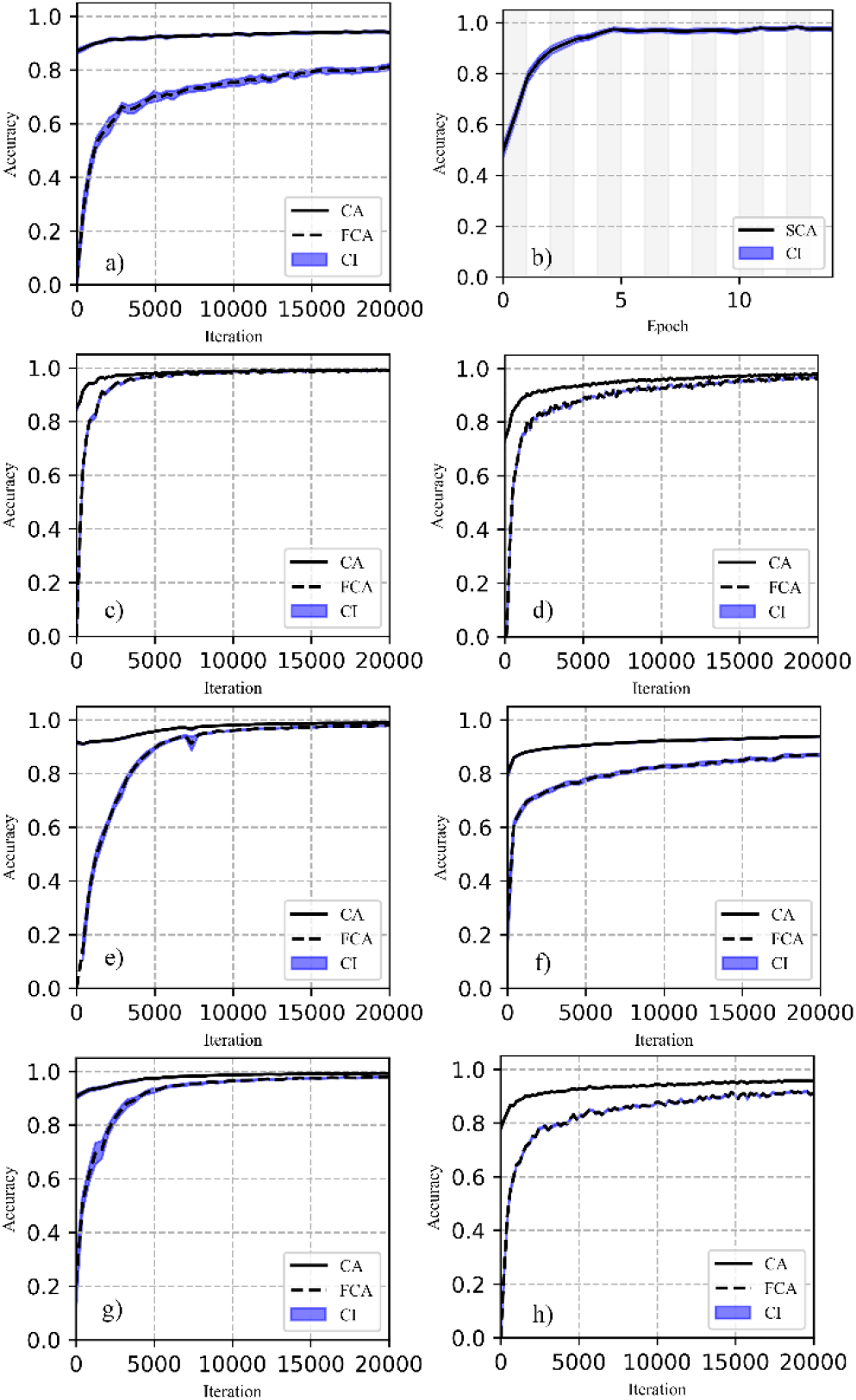
Classification accuracy (CA), foreground classification accuracy (FCA), confidence interval (CI), and scene classification accuracy (SCA) for full model (a), scene classification model (b), High concentration model (c), *Nocticula* model (d), *Phaeocystis* model (e), Pteropoda model (f), Shrimp model (g), and low-density model (h).

**Fig. 4.**
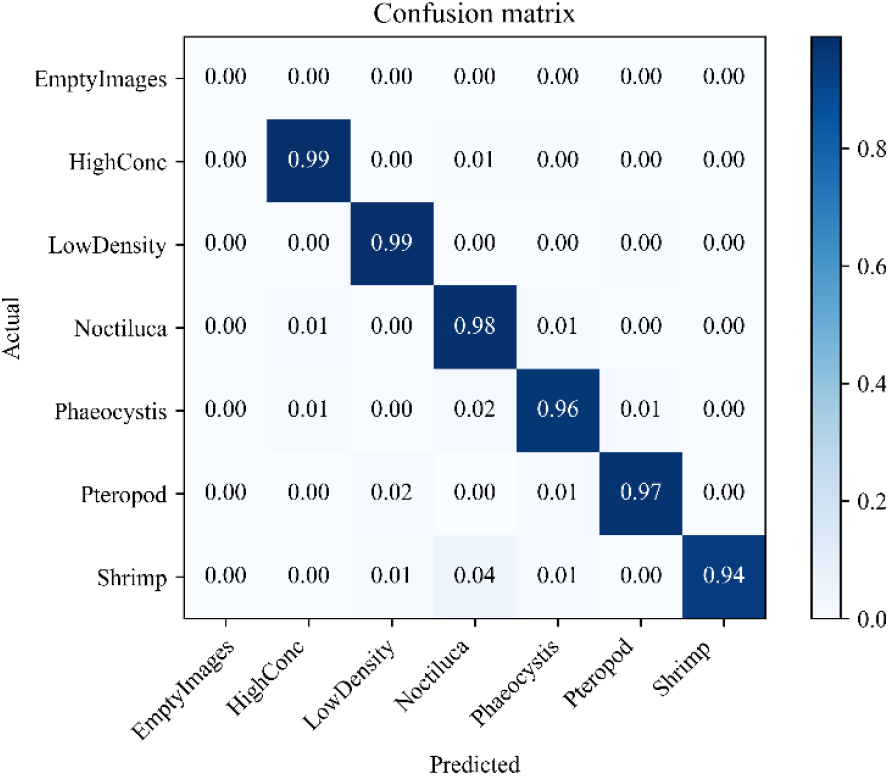
Normalized confusion matrix for the scene classification model with each row representing an actual class example and each column representing the state of a predicted class.

Both the full model and scene-specific model performed well on large organisms (Fig. 5,6; Table III). In the high-concentration scene, both procedures effectively identified and segmented targets from very noisy and low contrast backgrounds (Figs 5a, b). The scene-specific approach performed slightly better than the full model (Table III). In the shrimp scene, both procedures identified and segmented shrimp correctly, while the full model found more *Noctiluca* than the shrimp scene-specific model (Figs 5c, d, Table III); this disparity was caused by the larger sample of labelled *Noctiluca* in the full model library (4,950 individuals) than in the shrimp scene-specific library (782 individuals, Table I). In the low-density scene, both procedures performed well (Figs 5e, f) except that the shrimp scene-specific model tended to miss targets due to misclassification (Table III).

**Fig. 5.**
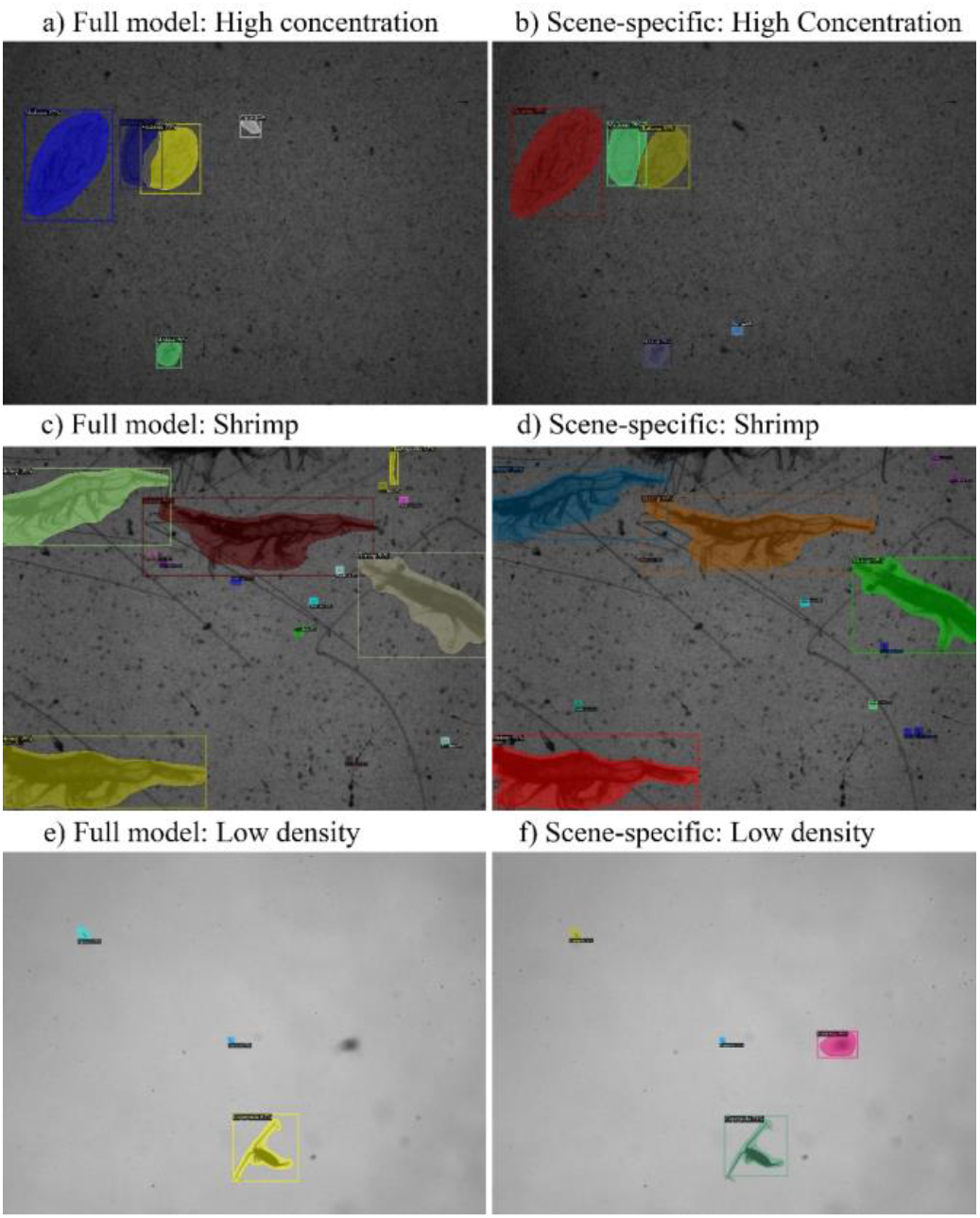
Examples of processed underwater plankton images from three different scene categories using the full model and scene-specific models.

**Fig. 6.**
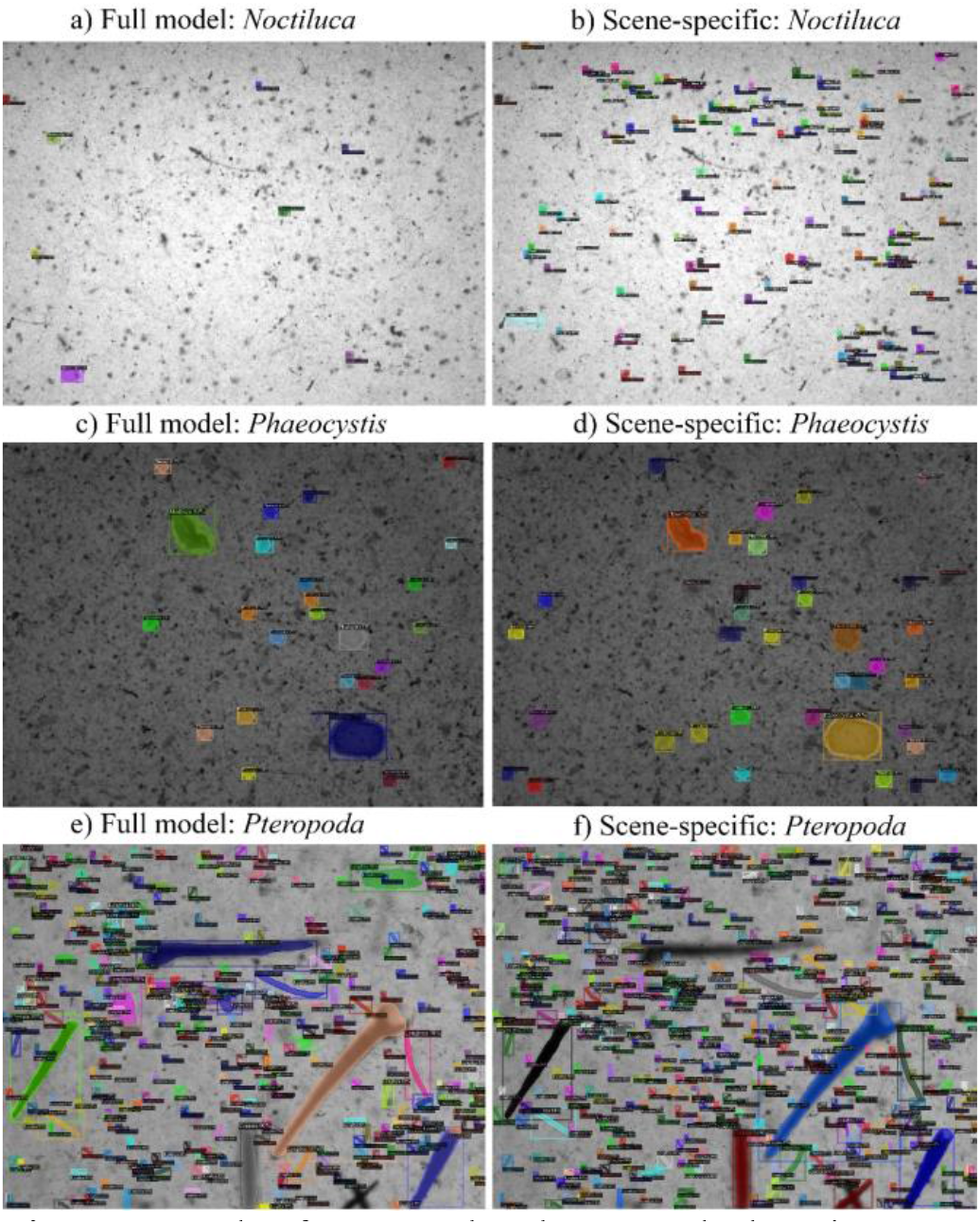
Example of processed underwater plankton images with high complexity.

**TABLE III.**
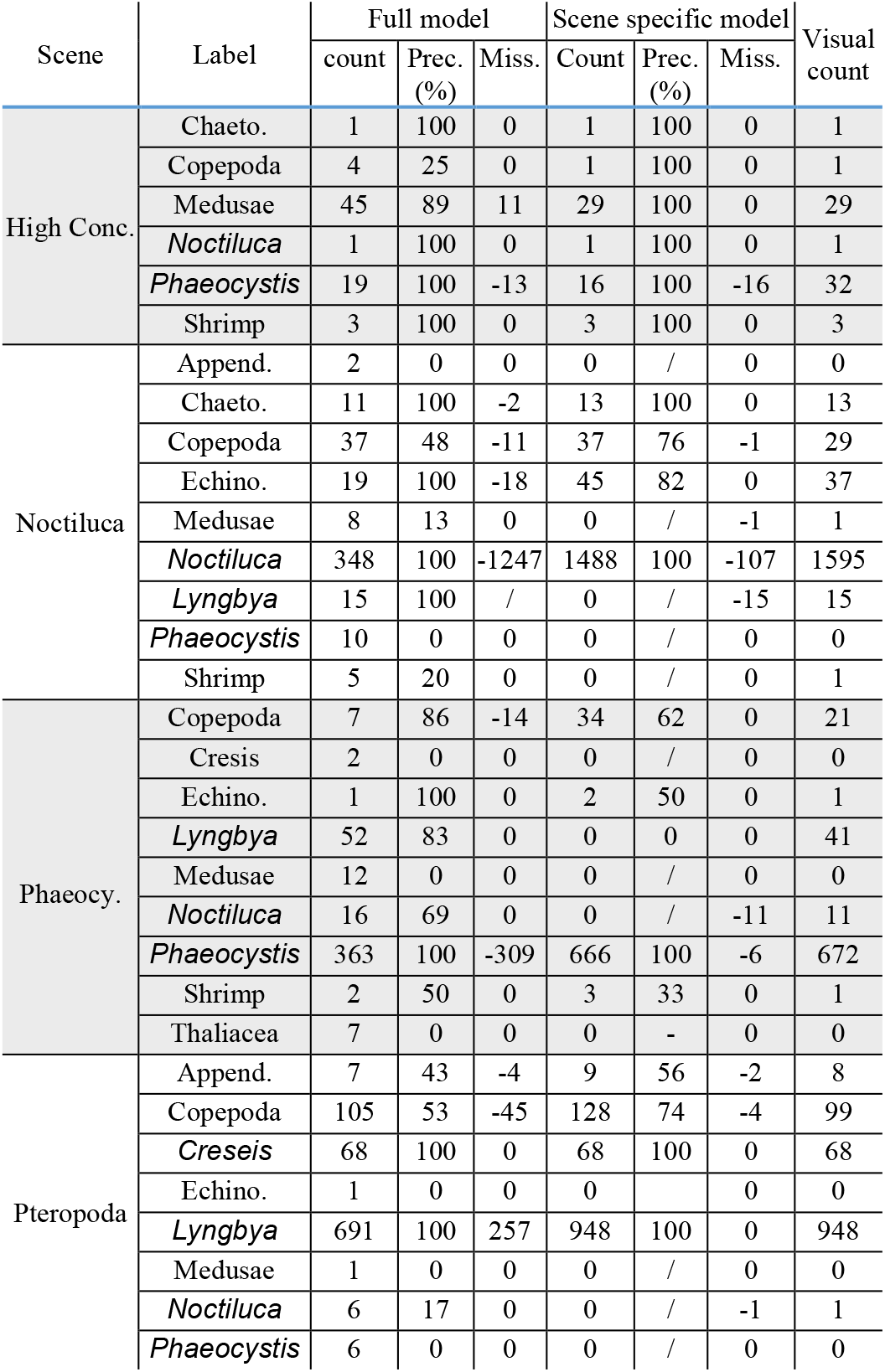
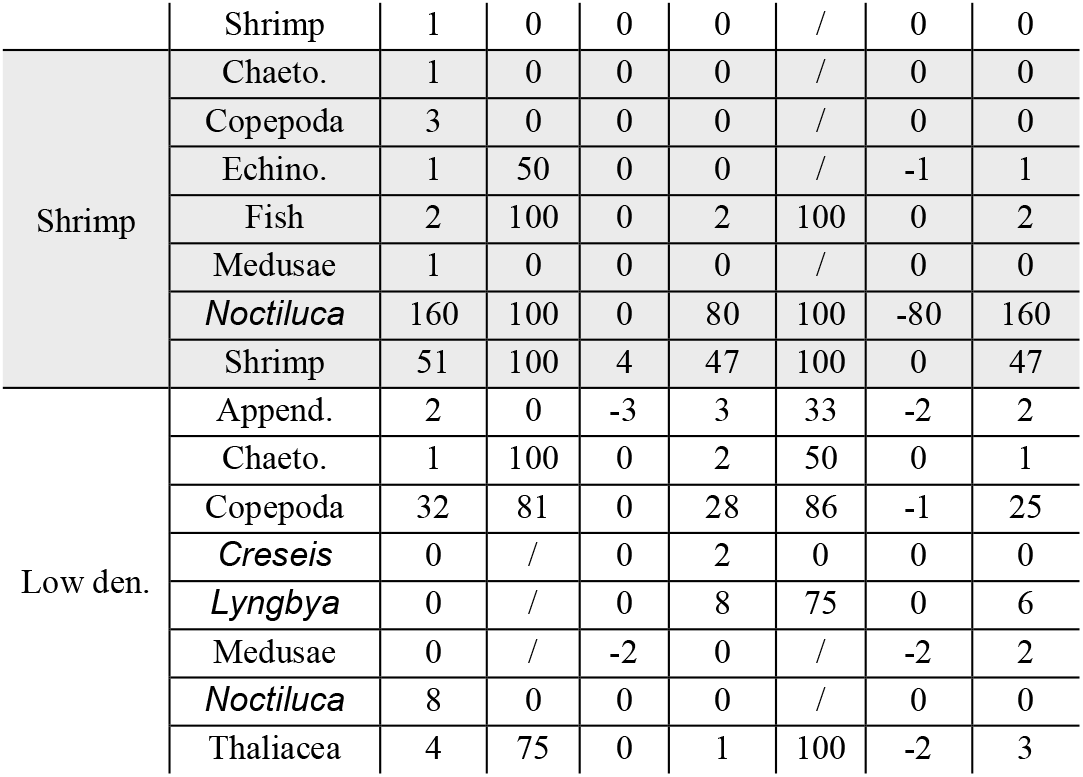
Results from the Processed Underwater Plankton Images Using the Full Model, Scene-specific Models, AND Visual Counts.^a,b^

When image complexity increased, the scene-specific models generally outperformed the full model with improved precision and reduced missing targets (Figs 6, Table III). For example, in the image from the *Noctiluca* scene (Fig. 6a, b), the full model only identified 5 *Noctiluca* individuals and 3 misclassified colonies of *Phaeocystis*, while the scene-specific model identified 130 *Noctiluca* individuals and 1 misclassified Enchinodermata larvae. In the 20 processed images, the full model missed 78% of *Noctiluca* individuals without considering misclassified individuals, despite the full model’s highly labelled *Noctiluca* library (∼35% individuals labelled). In the image from the *Phaeocystis* scene, the full model identified 24 colony stage of *Phaeocystis* and 2 misclassified medusae, while the *Phaeocystis* scene-specific model identified 38 colonies of *Phaeocystis* (Figs 6c, d). In the 20 processed images, the full model missed 46% of colonies,without considering misclassified individuals. In the image from the Pteropoda scene, the full model recorded 1 Copepoda, 5 Pteropoda, and 351 *Lyngbya*, while the scene-specific model recorded 4 Copepoda, 5 Pteropoda, and 505 *Lyngbya* (Figs 6e, f). In the 20 processed images, the full model missed ∼27% of *Lyngbya* individuals.

The accuracy of the scene-specific model for other small organisms, like copepods, is also much higher than the full model (Table III). The full inclusion model approach also tends to merge multiple RoIs for small organisms, such as *Lyngbya* clusters, which were often falsely recognized as a single *Lyngbya* individuals (Figs 6e, f). The occurrence of merged RoIs increased as we increased the number of large organisms, such as mysid shrimp, in the library. However, the scene-specific models effectively avoided the merging RoIs issue.

### B. General applications

Effectively extracting RoIs is a necessary, but challenging, first step in image recognition. Using benthic images as an example, it was difficult to extract intact RoIs from the background using traditional binarization methods, like adaptive thresholding, because the resulting RoIs were disconnected and fragmented into small pieces (Figs 7a, b). For images with aggregated benthic organisms (Fig 7c), the traditional binarization approach was even less effective and failed to yield any meaningful results. By comparison, the proposed approach performed well on low-contrast benthic images (Fig 7c, d). The number of images, scene categories, identified taxonomic groups, and labelled individuals for the full model and scene specific models are provided in Table 4. For the full model, the foreground classification accuracy and classification accuracy reached 99% and 95% respectively. For the scene-specific models, the accuracy of scene classification model reached 98%, the foreground classification accuracies ranged 97% -99%, and classification accuracies ranged 96% - 97% (Table 4). When we applied the trained models to the selected benthic images, the full model missed some individuals of the dominant taxonomic groups, ∼13% of sand dollar, ∼60% of brittle star, and ∼67% of sea anemone (Table 4). The scene-specific model often misclassified sea anemones as shells, and *vice versa*. Given the small number of labelled organisms in the library, we expect that the performance of both models to improve rapidly as the number of labelled organisms increases.

**Fig. 7.**
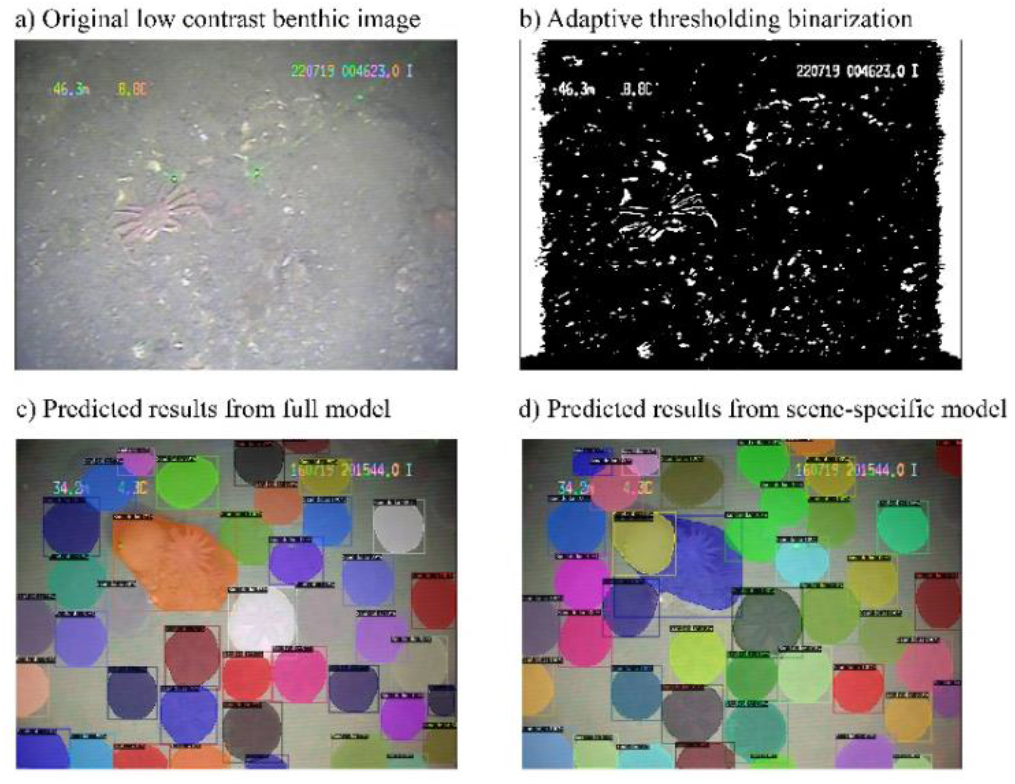
Benthic video images: a) an original benthic video image with a crab, b) binarized image using an adaptive thresholding approach to segment potential regions of interest, c) predicted results for an aggregated scene image using the full model, and d) predicted results for the same aggregated scene image using scene-specific model. Note in image c), there are a few sand dollars that were missed and a merge of a sand dollar and a sea anemone.

**TABLE IV.**
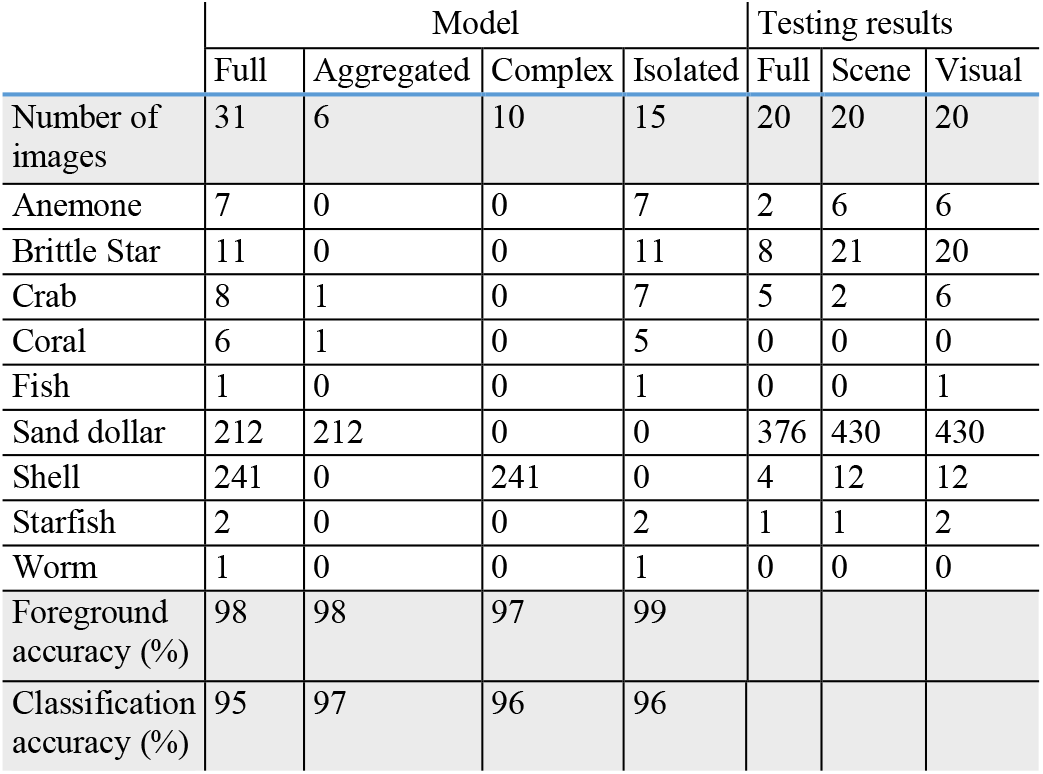
Number of Images, Identified Taxonomic Groups, Number of Labelled Individuals and Accuracies for the Full Model and Scene-specific Models.

Sonar images are often in relatively low resolution and lack details at individual level (Figs 8a, d). In many cases, soft-bodied jellyfish were obscured, with only the bell portion of their bodies visible. The number of images, scene categories, identified taxonomic groups, and labelled individuals are provided in Table V. For the full model, the foreground classification accuracy and classification accuracy reached 94% and 97% respectively. For the scene-specific models, scene classification accuracy was 96%, the foreground classification accuracies ranged from 92%-100%, and classification accuracies ranged from 90%-99% (Table V). The full model often had trouble distinguishing mysid swarms from small pelagic fish schools (Fig 8b, e); it tended to confuse small forage fish schools and mysid schools (Fig 8e). The scene-specific model successfully identified mysid swarms (Fig 8c) and enumerated small forage fish within the small pelagic fish school (Fig 8f). In summary, the full model overestimated the number of small fish, jellyfish and mysid swarms, while results from the scene-specific model were consistent with visual counts (Table V).

**Fig. 8.**
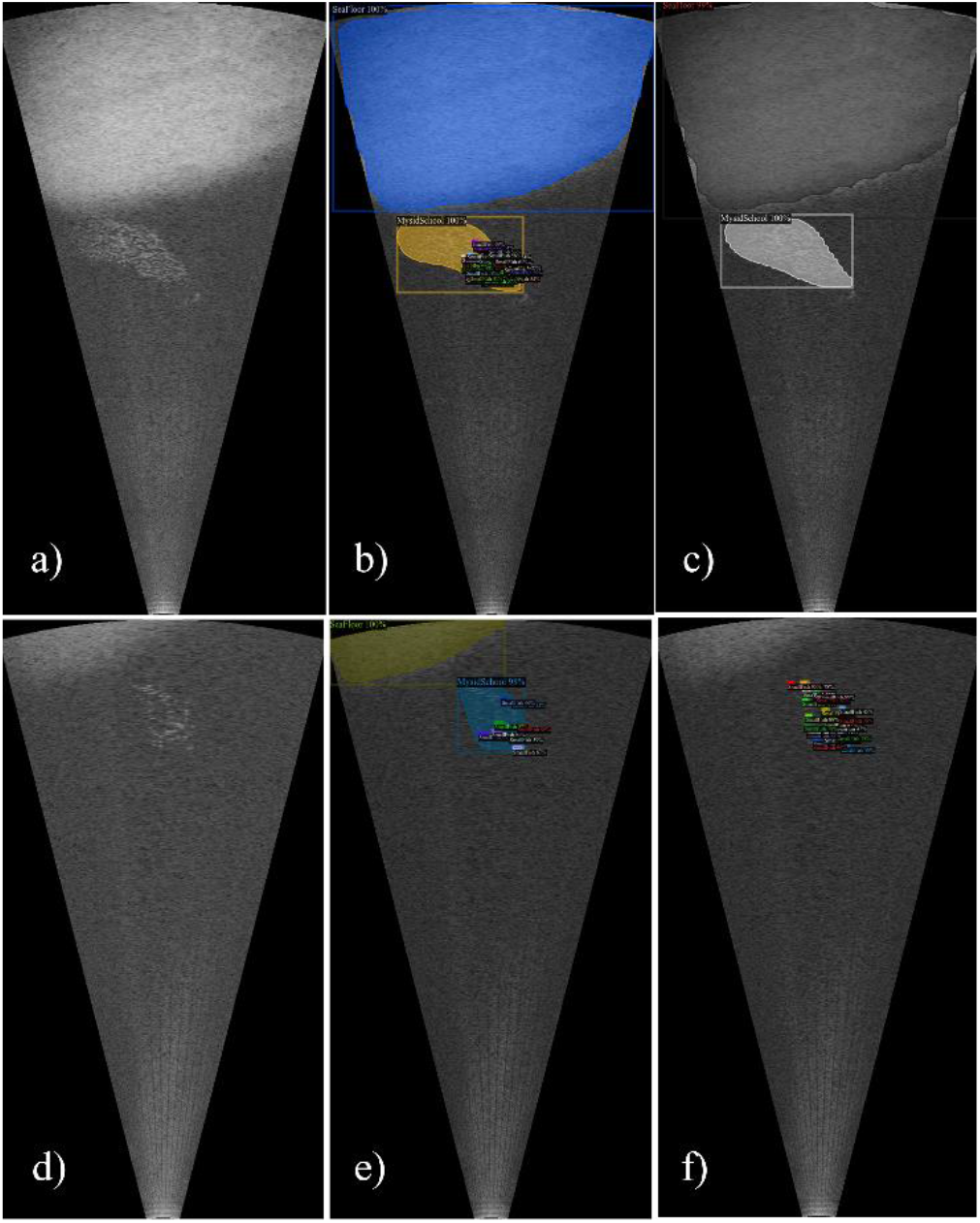
Sonar images: a) a sonar image showing seafloor and a swarm of mysids in the middle water column, and b) predicted results from the full model, c) predicted results from the scene-specific result; d) seafloor and a near bottom school of small forage fish, e) predicted results from the full model, and f) predicted results from the scene-specific model.

**TABLE V.** Number of Sonar Images, Identified Taxonomic Groups, Labelled Individuals and Accuracies for the Full Model AND Scene-specific Models.

|  | Number of images | Sea Floor | Small Fish | Mysid Swarm | Jellyfish | Foreground accuracy (%) | Class. accuracy (%) |
| --- | --- | --- | --- | --- | --- | --- | --- |
| Full model | 249 | 154 | 6346 | 159 | 130 | 94 | 97 |
| EmptyFloor | 30 | 30 | 0 | 0 | 1 | 100 | 99 |
| FishFloor | 29 | 29 | 1757 | 0 | 0 | 94 | 90 |
| FishNoFloor | 16 | 0 | 2949 | 0 | 8 | 92 | 91 |
| JellyfishFloor | 30 | 30 | 683 | 9 | 40 | 98 | 96 |
| JellyfishNoFloor | 29 | 0 | 613 | 2 | 55 | 100 | 98 |
| MysidFloor | 61 | 61 | 154 | 74 | 0 | 100 | 99 |
| MysidNoFloor | 51 | 0 | 0 | 74 | 26 | 100 | 99 |
| Full model | 20 | 17 | 634 | 5 | 6 |  |  |
| Scene model | 20 | 4 | 487 | 4 | 4 |  |  |
| Visual counts | 20 | 17 | 503 | 4 | 4 |  |  |

## IV. CONCLUSION

The new approach takes advantage of recent progress in artificial intelligence and is designed to address several challenges related to the distribution patterns of marine organisms in different environments. The inclusion of RPN significantly increased our ability to separate each RoI from its background. The RPN first generates a set of region proposals for each object, classifies each proposed region as foreground or background, and finally it produces the best fit region proposal for each object. The RPN model also identified non-target objects, and therefore could effectively reduce the number of objects that needed to be classified by the following CNN model. This initial step leads to subsequent increases in accuracy, reduced false positives, and reduced computational demands. Results suggest that the RPN approach is an ideal candidate as a common approach for object detection in different types of images, from marine biome.

When compared to a single, unbiased Mask R-CNN model for all images, the advantages of the proposed framework are threefold. First, a single unbiased classifier often oversamples rare groups resulting in more false positives and under-samples abundant groups leading to more false negatives. A scene-specific classification model for each scene takes into account the inherently patchy distribution of marine organisms meaning that dominant taxon could be different in different images. A scene-specific model for each scene also allows a better match in data distribution between samples and libraries, and therefore increases accuracy in both object extraction and recognition. Second, the scene specific approach provides a better method for dealing with large size differences among objects, for example, an algal cell versus an adult krill. A single unbiased classifier underestimates or completely misses small organisms. Third, the scene specific model reduces the variation and uncertainty among images by separating images with similar layout into the same scene which subsequently improves the model performance.

The Mask R-CNN approach also addresses the long-standing issue of broken and overlapping objects. Traditional binarization approaches often generate broken objects resulting in over segmentation which leads to more errors during classification as illustrated in the benthic images. The RPN used in the new approach has a clear advantage over other segmentation approaches, such as thresholding binarization. With advances in optics leading to significant increases of camera depth of field in order to increase the imaging volume, the likelihood of overlapping objects becomes a more pronounced issue. The new approach preserves the advantages of depth of field by computationally reducing the degree of overlap between objects (Fig 9). In the test image, there were 31 *Creseis acicula* and 5 of them were either out of focus or only partially presented. Several *C. acicula* were overlapping. The program accurately recognized most individuals and only over-segmented one individual, with a final count of 27 individuals. Finally, the new approach effectively eliminates false positives by implementing a simple binary classifier in the RoI segmentation step.

**Fig. 9.**
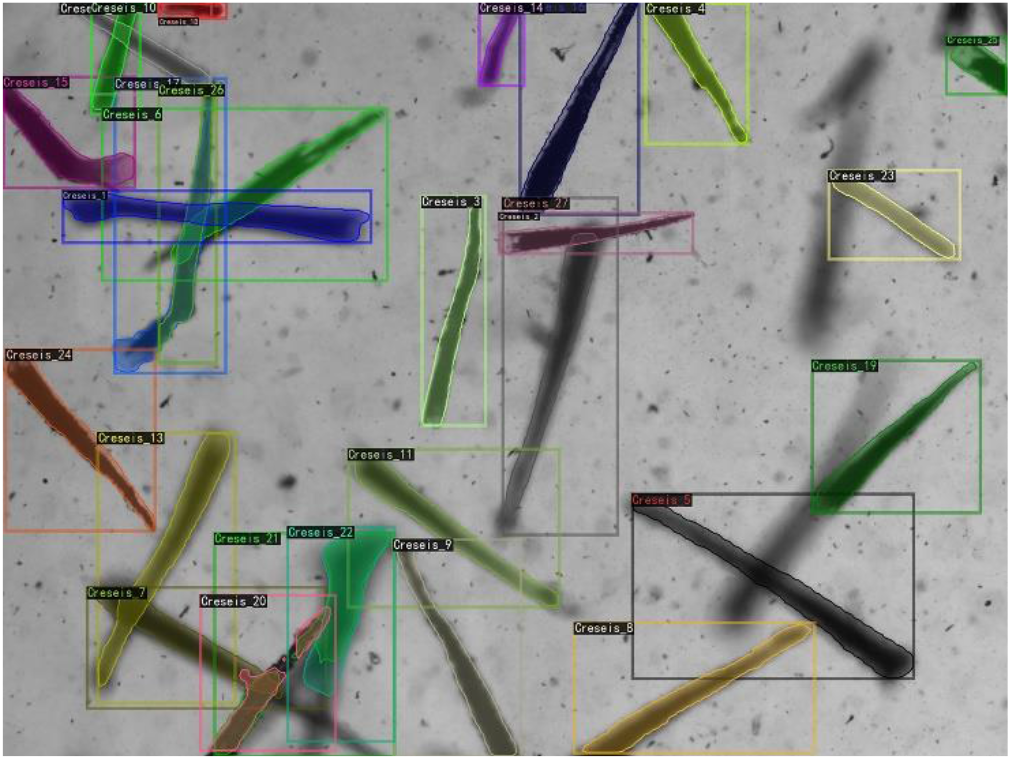
A plankton image dominated by *Creseis acicula* to illustrate the overlapping issue. In total, there were 30 individuals by visual examination and 5 were either partially presented or out of focus. The scene-specific model recognized 26 individuals with one over-segmented individual.

The proposed framework can process different types of images as illustrated in the present study: microscopic, complex plankton images, benthic video images, and sonar images. Furthermore, this unified framework fundamentally addresses the issue of customized system-specific algorithms and provides an opportunity to compare different systems within the same image processing framework. We conclude that the proposed framework can greatly facilitate the deployment of imaging systems in ecological studies by enabling rapid bulk image processing and extraction of useful ecological information.

## ACKNOWLEDGMENT

We thank Dr. Cooper for providing benthic videos. PlanktonScope images were collected by Bi and Ying in Shenzhen coastal waters, and sonar images were collected by Bi, Lankowicz, and Shahrestani in Chesapeake Bay.

## Footnotes

a Negative missing values indicate individuals that were not detected and misclassified; positive missing values indicate possible over-segmentation and dual labelled objects in which an object was recognized as more than one taxonomic group based on a 40% probability threshold.

b For small semi-transparent organisms like *Noctiluca* and *Lyngbya*, visual counts were performed based on the model output in which identified organisms were labelled.

## Notes

This work was supported by the National Key Research and Development Program of China under Grant 2017YFC1403600, and Major Scientific and Technological Innovation Project of the Shandong Provincial Key Research and Development Program under grant 2019JZZY020708

### Competing Interest Statement

The authors have declared no competing interest.

